# A bacterial artificial chromosome (BAC)-vectored noninfectious replicon of SARS-CoV-2

**DOI:** 10.1101/2020.09.11.294330

**Authors:** Yang Zhang, Wuhui Song, Shuiye Chen, Zhenghong Yuan, Zhigang Yi

**Affiliations:** Key Laboratory of Medical Molecular Virology (MOE/NHC/CAMS), School of Basic Medical Sciences, Shanghai Medical College, Fudan University, Shanghai, 200032, PR China; Shanghai public health clinical center, Fudan University, Shanghai, 201508, PR China

## Abstract

Vaccines and antiviral agents are in urgent need to stop the COVID-19 pandemic. To facilitate antiviral screening against SARS-CoV-2 without requirement for high biosafety level facility, we developed a bacterial artificial chromosome (BAC)-vectored replicon of SARS-CoV-2, nCoV-SH01 strain, in which secreted *Gaussia* luciferase (sGluc) was encoded in viral subgenomic mRNA as a reporter gene. The replicon was devoid of structural genes spike (S), membrane (M), and envelope (E). Upon transfection, the replicon RNA replicated in various cell lines, and was sensitive to interferon alpha (IFN-α), remdesivir, but was resistant to hepatitis C virus inhibitors daclatasvir and sofosbuvir. Replication of the replicon was also sensitive overexpression of zinc-finger antiviral protein (ZAP). We also constructed a four-plasmid *in-vitro* ligation system that is compatible with the BAC system, which makes it easy to introduce desired mutations into the assembly plasmids for *in-vitro* ligation. This replicon system would be helpful for performing antiviral screening and dissecting virus-host interactions.

## Introduction

The pandemic COVID-19 has infected over 26 million people and caused over 800,000 mortalities. (https://www.who.int/emergencies/diseases/novel-coronavirus-2019). It is caused by infection with a novel *beta* coronavirus SARS-CoV-2 ^1–3^. Vaccines and antiviral agents are in urgent need to stop the pandemic. Despite great progresses on SARS-CoV-2 vaccine development and clinical trails ^4^, the protection efficacy of the vaccines still remains to be determined. There have been trials of antiviral agents such as remdesivir and chloroquine for COVID-19 treatment, however, efficacy of these antiviral agents remains uncertainty ^5–7^. Development of convenient tools for antiviral screening will speed up seeking effective antiviral agents against SARS-CoV-2. Recently, infectious clones of SARS-CoV-2 with reporter genes ^8, 9^ provide elegant tools for antiviral development. However, due to the safety issue and requirement for biosafety level 3 laboratory, usage of these infectious clones is limited. Non-infectious replicon system that recapitulates authentic viral replication without virion production can be used to perform screening for antivirals that target viral replication process.

SARS-CoV-2 contains an approximate 29kb, single stranded, positive sense RNA genome. About two-thirds of the viral genome encodes open reading frames (ORFs) for translation of the replicase and transcriptase proteins, the only ORFs translated from the viral genome. The translated replicase and transcriptase proteins engage viral genome to assemble the replicase-transcriptase complex on endoplasmic reticulum membrane, forming a membranous compartment. Within the membranous compartment, replicase-transcriptase complex initiates viral replication and transcription. Transcription of the 3'-most third genomes by viral replicase-transcriptase generates various subgenomic mRNAs that encode structural proteins and accessory genes ^10^. Structural proteins include the spike (S), membrane (M), envelop (E) proteins and nucleocapsid (N) participate in virion assembly ^10^. In this study, we generated a replicon system for SARS-CoV-2, nCoV-SH01 strain with secreted *Gaussia* luciferase (sGluc) as a reporter gene. The cDNA of viral genome with deletion of S, M, E genes was cloned into a bacterial artificial chromosome (BAC) vector. The reporter gene sGluc was encoded in subgenomic viral RNA. The viral RNA was transcribed *in vitro* by T7 polymerase. Upon transfection into cells, the viral replication was detected, as evidenced by expression of subgenomic viral RNA-encoded sGluc. The viral replication was sensitive to interferon alpha (IFN-α), remdesivir, but was resistant to hepatitis C virus inhibitors daclatasvir and sofosbuvir. The replicon genomes could also be assembled by *in-vitro*-ligation of four DNA fragments and the RNA generated by the *in-vitro*-ligated DNA template was capable of replication as the RNAs derived from the BAC-template. Thus, we provided a simple SARS-CoV-2 replicon system for antiviral development.

## Results

### Construction of a bacterial artificial chromosome (BAC) based SARS-CoV-2 replicon

Total RNAs were extracted from SARS-CoV-2 (nCoV-SH01) infected cells ^11^, and then reversely transcribed by superscript IV with random primer. Totally, 20 fragments with approximate l.5kb-length encompassing the whole viral genomes were amplified with specific primers according to the illumina-sequenced viral genome (MT121215), cloned and sequenced. The fragments were then assembled by fusion PCR and subcloning into larger fragments A (1-8586nt), B (8587-15102nt), C (15103-21562nt) and D and cloned into a homemade cloning vector pLC-Zero-blunt (Fig.1). Took a similar strategy for construction of SARS-CoV replicon ^12^, we deleted the structural protein genes and retained the N gene and essential promoter regions. We replaced the S gene region with a reporter gene cassette, including secreted *Gaussia* luciferase (sGluc), foot-and-mouth disease virus (FMDV) 2A peptide and blasticidin (BSD), whose expression was driven by the promoter of S gene in the subgenomic mRNA (Fig. 1). To facilitate cloning, a *BamHI* site was introduced downstream the genome position of 21562 (nt) in the pLC-nCoV-C plasmid. T7 promoter was added before the 5′ viral genome in the fragment A for *in vitro* transcription with T7 polymerse. The 3′ viral genome was flanked with polyA30, hepatitis delta virus ribozymes (HDVr) and terminator sequence for T7 polymerase (T7T) (Fig. 1), which facilitates *in-vitro* transcription without linearization and production of precise polyadenylated viral RNA. The fragments of A, B, C and D were assembled further into AB and CD in pLC-Zero-blunt and then sequentially cloned into a modified BAC vector to get the final plasmid pBAC-sgnCoV-sGluc (Fig. 1).

**Fig. 1.**
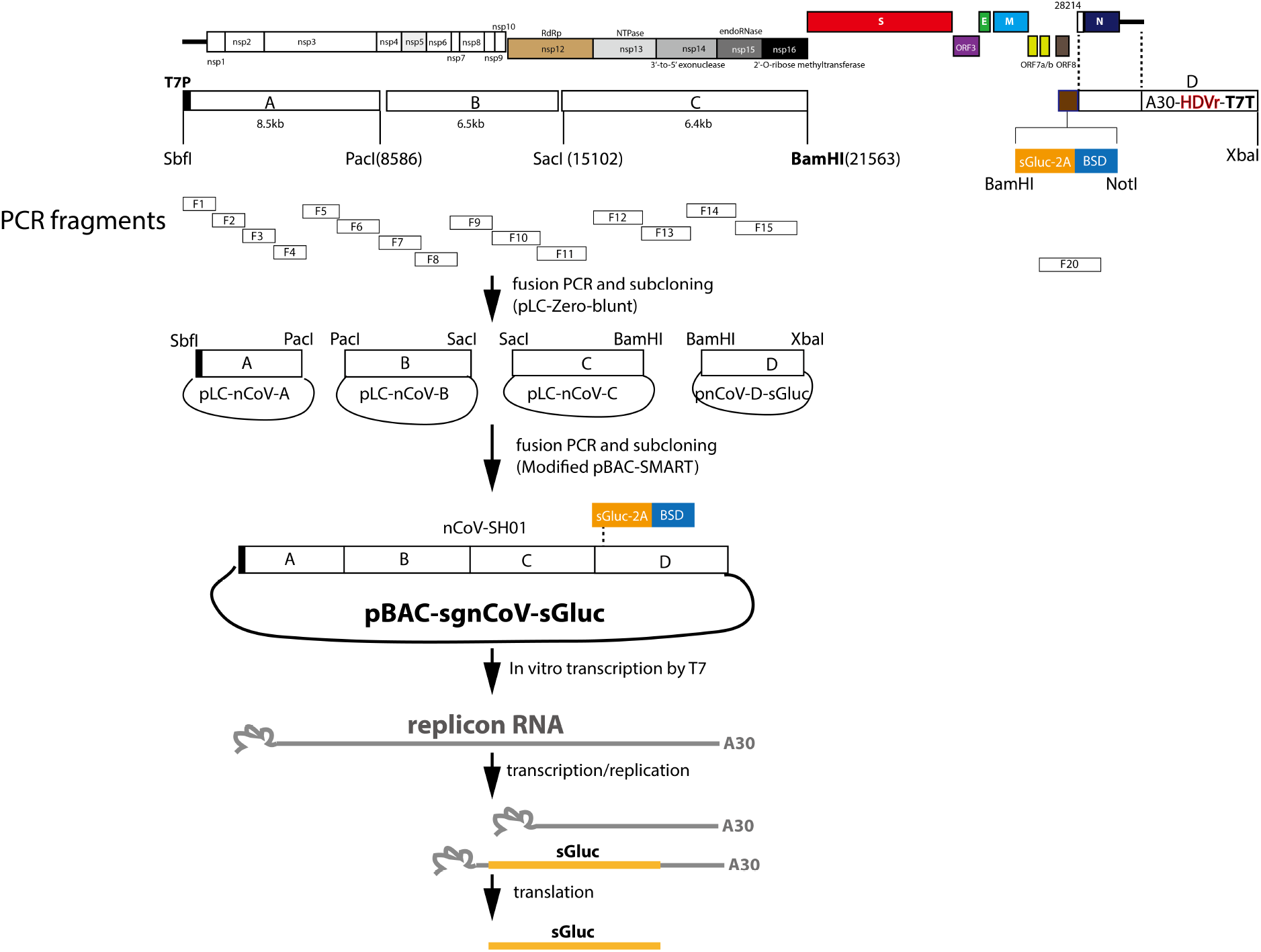
Schematic of construction of BAC-based replicon of SARS-CoV-2. Twenty fragments encompassing the whole viral genomes were amplified, cloned and sequenced. Four larger fragments A (1-8586nt), B (8587-15102nt, C (15103-21562nt) and D with deletion of structural protein genes and addition of reporter gene cassette were assembled and cloned. To facilitate cloning, a BamHI site (**in bold**) was introduced downstream the genome position of 21562 (nt). In fragment A, T7 promoter (T7P) was added. In fragment D, an expression cassette containing secreted *Gaussia* luciferase (sGluc), foot-and-mouth disease virus (FMDV) 2A peptide and blasticidin (BSD) was added. The 3′ viral genome was flanked with polyA30, hepatitis delta virus ribozymes (HDVr) and terminator sequence for T7 polymerase (T7T). Then the fragments were assembled and sequentially cloned into a modified BAC plasmid. Upon transfected into cells, the replicon RNA can be used as template for RNA replication or transcription to produce subgenomic RNA. The sGluc subgenomic RNA is translated to produce sGluc.

### Replication of SARS-CoV-2 replicon in cells

The plasmid pBAC-sgnCoV-sGluc was used directly as template for *in-vitro* transcription to produce 5′-capped replicon RNA. Replicon RNA was then co-transfected with N mRNA into various cell lines. RNA replication was monitored by measuring the secreted *Gaussia* luciferase activity in the supernatants. Enzymatic dead mutants (759-SAA-761) of the RNA dependent RNA polymerase nsp12^13^ were introduced and the mutated replicon served as a non-replication control. As expected, SAA RNA did not replicate, without increase of luciferase activity in the transfected Huh7, Huh7.5, Vero and BHK-21 cells. In contrast, transfection of wild type (WT) replicon RNA resulted in obvious increase of luciferase activity (Fig. 2a, b, c, d), indicating active viral replication. Huh7.5 cell is a subclone of Huh7 cells with deficiency in RIG-I and MDA-5 signaling^14, 15^. Vero cell is routinely used for SARS-CoV-2 isolation. Notably, replicon replication was less efficient in Huh7.5 and Vero cells (Fig. 2b and c) whereas robust in BHK-21 cells (Fig. 2d), suggesting that cellular environment may regulate SARS-CoV-2 replication. In consistent with previous studies, co-transfection of N mRNA enhanced viral replication (Fig. 2a, b, c, d), which is probably due to the suppression of innate immune response^16^. For convenience, we tried to establish an N-expressed cell line (Fig.2e). Compared with GFP-expressed cells, Huh7 cells expressing N supported more robust viral replication (Fig. 2f).

**Fig. 2.**
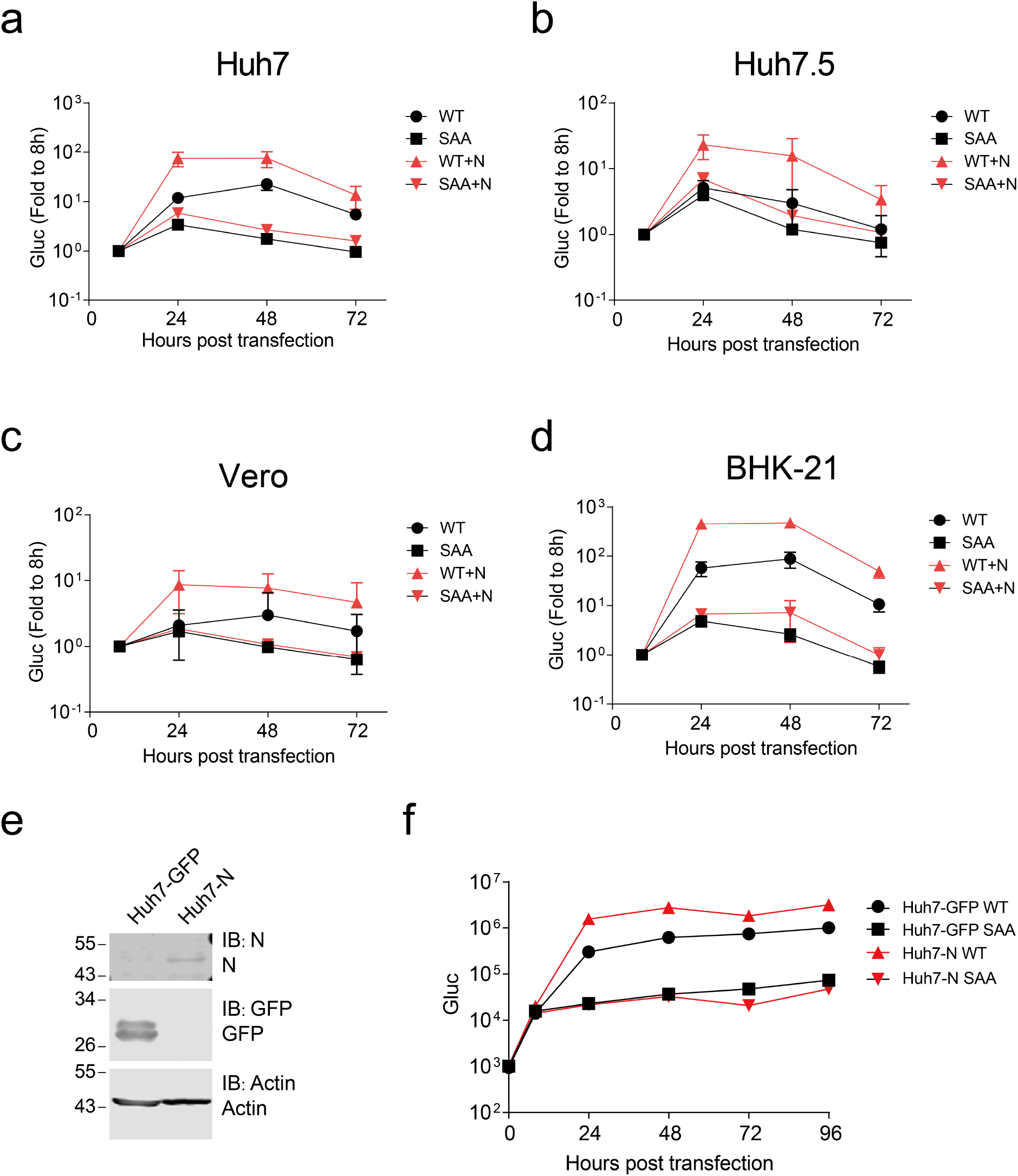
Replication of sgnCoV-sGluc in different cells. **a-d** Huh7, Huh7.5, Vero and BHK-21 cells were transfected with *in-vitro*-transcribed replicon RNA(WT) or the nsp12 polymerase active-site mutant (SAA). An mRNA encoding the SARS-CoV-2 N protein was co-transfected or not. The luciferase activity in the supernatants was measured at the time points indicated. Medium was changed at each time point. Data are shown as mean±SD (n=3). **e-f** Replication of replicon RNA in Huh7 cells overexpressed N protein. **e** Huh7 cells overexpressed GFP protein or N protein were analyzed by Western blotting with the indicated antibodies. **f** Huh7-GFP and Huh7-N cells were transfected with replicon RNA (WT or SAA). The luciferase activity in the supernatants was measured at the time points indicated. Medium was changed at 8 hours post transfection. Data are shown as mean±SD (n=3).

### Sensitivity of the SARS-CoV-2 replicon to antiviral agents

We tested the sensitivity of SARS-CoV-2 replicon to remdesivir, which has been demonstrated to inhibit SARS-CoV-2 viral infection^17^. Huh7 cells were first treated with remdesivir at various concentrations (10μm, 3.7μm, 1μm, 100nm, 10nm) as reported^17^, and then the cells were co-transfected with replicon RNA (WT or SAA) and N mRNA. The luciferase activity in the supernatants was measured at various time points after transfection. We found that at all concentrations, remdesivir effectively inhibited replicon replication to a similar level as SAA (Fig.3a). We also examined if the remdesivir inhibited established viral RNA replication. We first transfected replicon RNA, and then added remdesivir eight hours post transfection and monitored viral replication at various time points post treatment. Under this condition, remdesivir also reduced whereas did not completely block viral replication (Fig. 3b). Then we tested the sensitivity of SARS-CoV-2 replicon to other antiviral agents. Huh7 cells were first treated with interferon ahpha (IFN-α)(100U/ml), remdesivir (10 nM), daclatasvir (1 μM), sofosbuvir (10 μM) and 2’-C-Methylcytidine (2CMC)(50 μM) for four hours, then the cells were co-transfected with replicon RNA (WT or SAA) and N mRNA. The luciferase activity in the supernatants was measured at the various time points after transfection. IFN-α and remdesivir have been demonstrated to inhibit SARS-CoV-2 viral infection^17, 18^. Daclatasvir and sofosbuvir are direct antivirals targeting hepatitis C virus NS5A^19^ and RNA dependent RNA polymerase^20^, respectively. 2’-C-Methylcytidine is a nucleoside inhibitor of HCV NS5B polymerase ^21^. As shown in Figure 3c, IFN-α and remdesivir effectively inhibited sgnCoV-sGluc replication. Notably, IFN-α started to reduce the reporter gene expression at early time point (8 hours post transfection), manifested as lower luciferase activity then the SAA mutant, which suggests that IFN-α may block translation of the viral subgenomic mRNA. Remdesivir effectively inhibited the luciferase expression to a similar level of SAA. In contrast, sofosbuvir and 2’-C-Methylcytidine hardly reduced luciferase expression and daclatasvir had no effect on luciferase expression (Fig. 3c). These results demonstrate that SARS-CoV-2 replicon is sensitive to antiviral agents against SARS-CoV-2.

**Fig. 3.**
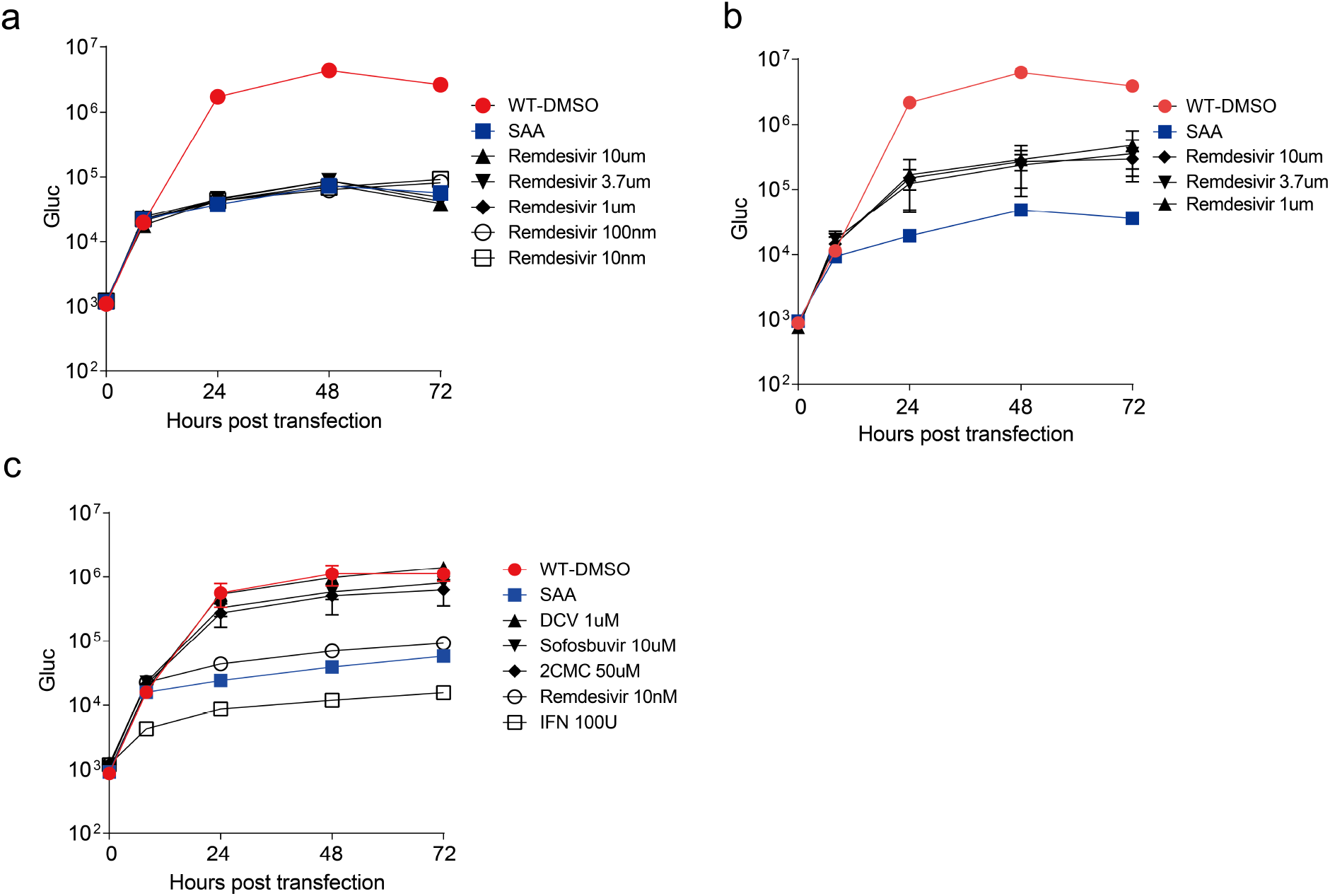
Sensitivity of the SARS-CoV-2 replicon to antiviral agents. **a** Huh7 cells were treated with remdesivir as the indicated concentration. Four hours later, cells were co-transfected with replicon RNA (WT or SAA) and N mRNA. The luciferase activity in the supernatants was measured at the time points indicated. **b** Huh7 cells were co-transfected with replicon RNA (WT or SAA) and N mRNA. Eight hours later, medium was changed with remdesivir as the indicated concentration. The luciferase activity in the supernatants was measured at the time points indicated. **c** Huh7 cells were treated with remdesivir (10nM), IFN-α(100 U/ml), daclatasvir(1 μM), sofosbuvir(10 μM), 2CMC(50 μM). Four hours later, cells were co-transfected with replicon RNA (WT or SAA) and N mRNA. The luciferase activity in the supernatants was measured at the time points indicated. Medium was changed at 8 hours post transfection. Data are shown as mean±SD (n=3).

### Sensitivity of SARS-CoV-2 replicon to overexpression of Zinc-finger antiviral protein (ZAP)

Zinc-finger antiviral protein recognizes CpG dinucleotide on non-self RNA and exerts antiviral activity^22^. There is extreme low CpG content of SARS-CoV-2 genome, suggesting SARS-CoV-2 may evolve under the pressure of ZAP^23, 24^. We generated a stable Huh7 cell line expressing the long isoform of ZAP (ZAPL) and examined the replicon RNA replication in the Huh7-ZAPL cells (Fig. 4a). There was about 10-fold reduction of replicon replication in Huh7-ZAPL comparing with that in GFP expressing cells (Huh-GFP) (Fig. 4b), suggesting replicon replication is sensitive to ZAPL overexpression.

**Fig. 4.**
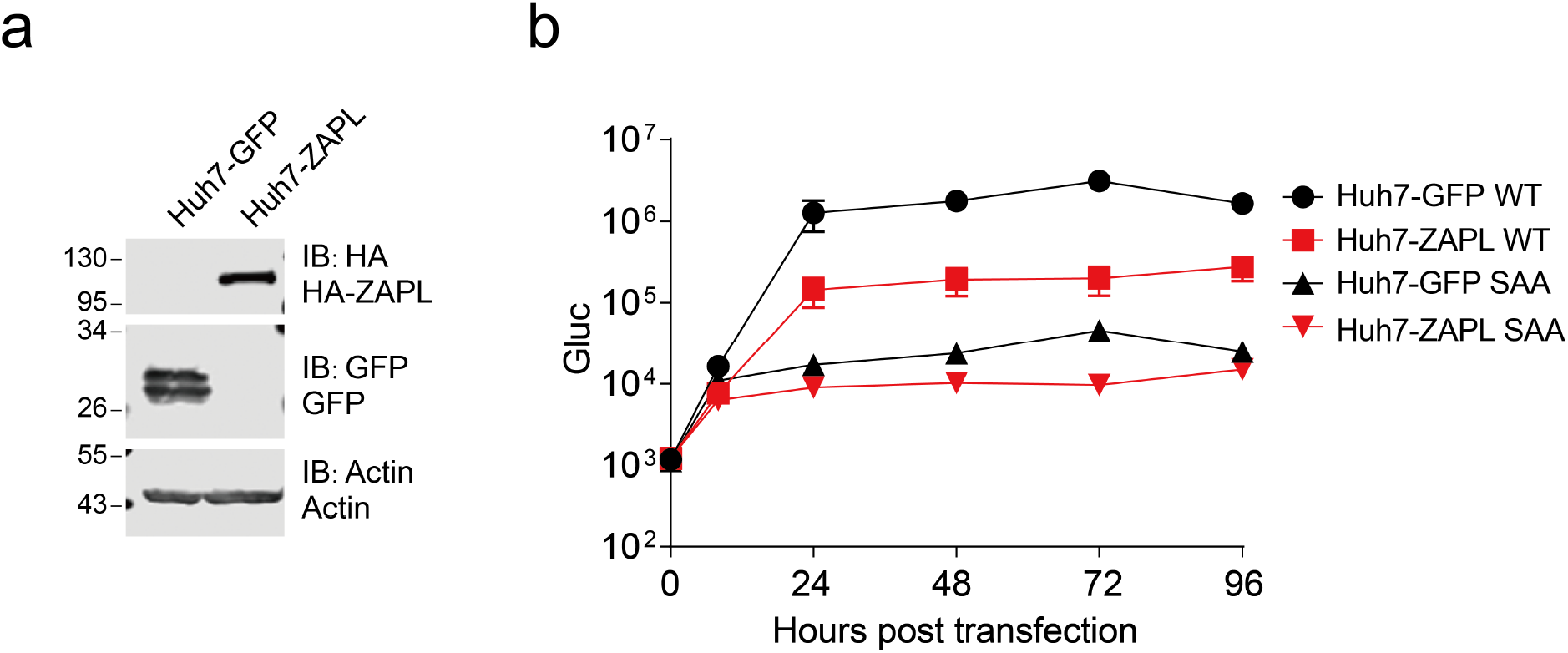
Sensitivity of SARS-CoV-2 replicon to overexpression of Zinc-finger antiviral protein (ZAP). **a** Huh7 cells overexpressed ZAPL protein or GFP protein were analyzed by Western blotting with the indicated antibodies. **b** Huh7-GFP and Huh7-ZAPL cells were co-transfected with replicon RNA (WT or SAA) and N mRNA. The luciferase activity in the supernatants was measured at the time points indicated. Medium was changed at 8 hours post transfection. Data are shown as mean±SD (n=3).

### Assembly of SARS-CoV-2 replicon by *in vitro* ligation

As the difficulties to manipulate with BAC vectors, we tried to assemble the four fragments A, B, C and D by *in-vitro* ligation. We introduced additional BsaI sites into the 5′ and 3′ of each fragment in the assembly plasmids pLC-nCoV-A, pLC-nCoV-B, pLC-nCoV-C, pnCoV-sGluc, retaining all the original restrictions enzymes. The fragments were released from the plasmid by BsaI digestion, and assembled by *in-vitro* ligation with T4 ligase (Fig. 5a). RNAs transcribed from the *in-vitro* ligated template replicated similar as the RNAs transcribed from BAC vector (Fig. 5b).

**Fig. 5.**
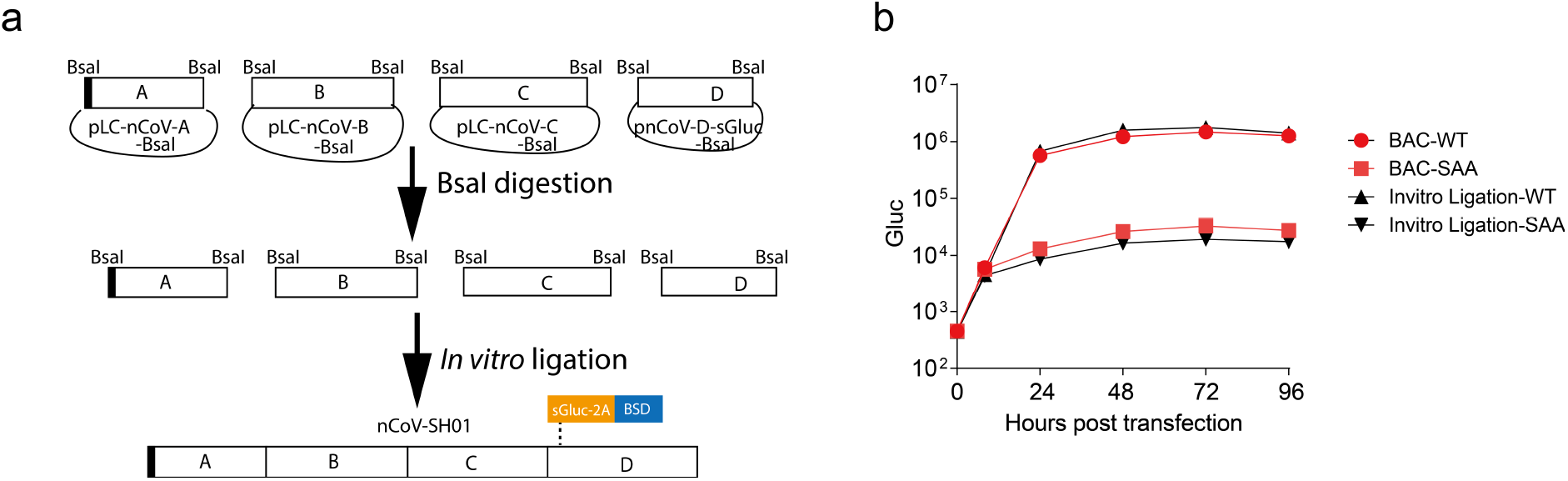
Assembly of SARS-CoV-2 replicon by *in vitro* ligation. **a** Schematic of the *in-vitro* ligation system for SARS-CoV-2 replicon. **b** Huh7 cells were co-transfected with replicon RNA (WT or SAA) and N mRNA generated by BAC-based system or *in-vitro* ligation system. The luciferase activity in the supernatants was measured at the time points indicated. Medium was changed at 8 hours post transfection. Data are shown as mean±SD (n=3).

## Discussion

In this study, we described a replicon system of SARS-CoV-2. In the replicon, we deleted the spike (S), membrane (M), envelop (E) genes that are essential for virion production, making it non-infectious and safe (Fig. 1). Upon transfection into various cells, the replicon RNA could replicate, manifested by the expression of subgenomic mRNA encoded sGluc (Fig. 2). The viral replication was inhibited by anti-SARS-CoV-2 antiviral agent remdesivir and by IFN-α but was not by antivirals against hepatitis C virus (Fig. 3). This replicon system avoids requirement for specific biosafety facilities. The BAC-vectored replicon system does not need *in vitro* ligation or recombination in yeast, which simplifies the experiment processes. Thus this replicon system would be used conveniently to perform antiviral screening against SARS-CoV-2. We also constructed a four-plasmid *in-vitro* ligation system that is compatible with the BAC system. Replicon RNAs produced from the *in-vitro* ligated replicate similarly with the RNAs transcribed from BAC plasmids (Fig. 5). It is easy to introduce desired mutations into the assembly plasmids for *in-vitro* ligation, which make it suitable for dissecting the effect of emerging mutations on viral replication and molecular mechanisms of viral replication.

## Material and methods

### Cloning

Total RNAs were extracted from SARS-CoV-2 (nCoV-SH01) infected cells ^11^, reversely transcribed by superscript IV (Invitrogen) with random primer. Totally 20 fragments with approximate l.5kb-length encompassing the whole viral genomes were amplified with specific primers according to the illumina-sequenced viral genome (MT121215), cloned into a homemade cloning vector pZero-blunt and sequenced. Four larger fragments A (1-8586nt), B (8587-15102nt, C (15103-21562nt) and D with deletion of structural protein genes and addition of reporter gene cassette were assembled by fusion PCR and subcloning, and then cloned into a homemade cloning vector pLC-Zero-blunt and pcDNA3.1 (invitrogen), respectively, resulted in plasmids pLC-nCoV-A, pLC-nCoV-B, pLC-nCoV-C, and pnCoV-D-sGluc. To facilitate cloning, a *BamHI* site was introduced downstream the genome position of 21562 (nt) in the plasmid pLC-nCoV-C. In fragment A, T7 promoter was added before the 5′ viral genome. In fragment D, a expression cassette containing secreted *Gaussia* luciferase (sGluc), foot-and-mouth disease virus (FMDV) 2A peptide (*NFDLL KLAGD VESNP GP*) and blasticidin (BSD) was added upstream the 5′-postion of viral genome. The 3′ viral genome was flanked with polyA30, hepatitis delta virus ribozymes (HDVr) and terminator sequence for T7 polymerase (T7T). Inactive mutants (759-SAA-761) of the RNA dependent RNA polymerase nsp12 was introduced into the C fragment at the predicted catalytic residues (759-SDD-761)^13^ by fusion PCR mediated mutagenesis.

To assemble the four fragments into bacterial artificial chromosome (BAC) vector, first we modified the pSMART-BAC v2.0 (Lucigen) to get ride of unwanted restriction enzymes and added *AatII and XhoI* sites to facilitate cloning by multiple rounds of fusion-PCR mediated mutagenesis. The fragments were then sequentially cloned into the BAC vector. We first assemble the fragment A and B, C and D by enzyme digestion to get the plasmid pLC-nCoV-AB and pLC-nCoV-CD, respectively. Then the AB fragments were cloned into the SbfI/XhoI site to generate pBAC-sgnCoV-AB. Then the CD fragments were ligated into the SacI/AsisI site to get pBAC-sgnCoV-sGluc. BAC plasmid was delivered into BAC-Optimized Replicator v2.0 Electrocompetent Cells (Lucigen) by electroporation and bacteria was propagated according to the manufacturer’s guide. Colonies were picked and cultured in LB medium containing 12.5 μg/ml chloramphenicol. L-arabinose was added to cultures when the OD_600_ reaches 0.2-0.3 to increase the plasmid copy numbers.

For assembly of SARS-CoV-2 replicon by *in-vitro*-ligation, we first got rid of the BsaI site on fragment C by fusion-PCR mediated synonymous mutagenesis. The BsaI sites were added into the 5′ and 3′ of the fragment A, B, C and D in the plasmids of pLC-nCoV-A, pLC-nCoV-B, pLC-nCoV-C, and pnCoV-D-sGluc by fusion PCR-mediated cloning, resulted in the plasmids pLC-nCoV-A-BsaI, pLC-nCoV-B-BsaI, pLC-nCoV-C-BsaI, and pnCoV-D-sGluc-BsaI. The plasmids retained all the original enzymatic sites, for convenience to swap into the BAC vector if desired. The fragments were released from the plasmids by *BsaI* digestion, after gel purification and ligated by T4 ligase.

To construct lentiviral vector expression plasmids, sequences encoding the GFP, SARS-CoV-2 nucleocapsid protein (N) were cloned into the XbaI/BsrGI site of pTRIP-IRES-BSD. Sequence encoding the long isoform of Zinc-finger antiviral protein (ZAPL) were synthesized by Wuxi Qinglan Biotech (Wuxi, China) and cloned into the XbaI/BamHI site of pTRIP-IRES-BSD. An HA tag was added into the N-terminal of ZAP. For production of N mRNA, sequence encoding N was first cloned into the KpnI/BamHI sites of phCMV to get the plasmid phCMV-N. All the plasmids were verified by Sanger sequencing. The detail information was available upon request.

### Cell lines

The human hepatoma cells Huh 7, baby hamster kidney cells BKH-21, Vero E6 cells were purchased from the Cell Bank of the Chinese Academy of Sciences (www.cellbank.org.cn) and routinely maintained in Dulbecco’s modified medium supplemented with 10 % FBS (Gibco) and 25 mM HEPES (Gibco). Huh 7.5 (Kindly provided by C. Rice) cells were routinely maintained in a similar medium supplemented with non-essential amino acids (Gibco). Huh7-GFP, Huh7-N, Huh7-ZAPL cell line was routinely maintained in the medium supplemented with 0.5 μg/ml blasticidin.

### Lentivirus pseudoparticle

VSV-G-pseudotyped lentiviral particles were prepared by co-transfection of VSV-G, HIV gag-pol and lentiviral provirus plasmids into HEK293T cells. The medium overlying the cells was harvested at 48 h after transfection, filtered through a 0.45-μm filter, and stored at −80°C. Cells were transduced with the pseudoparticles in the presence of 8 μg/ml Polybrene.

### Inhibitors

Remdesivir (GS-5734), Daclatasvir (S1482), Sofosbuvir (GS-7977) were purchased from Selleckchem, 2’-C-Methylcytidine (HY-10468) was purchased from MedChem Express, IFN-α (11200-2) was purchased from PBL.

### Antibodies

Anti-β-actin antibody (Sigma; A1978) was used at 1:5000 dilution; Anti-HA antibody (CST; 37243) was used at 1:000 dilution; Anti-GFP antibody (Santa Cruz;sc-9996) was used at 1:1000 dilution; Anti-ZAPL antibody (Proteintech; 16820-1-AP) was used at 1:1000 dilution; Anti-N antibody(GeneTex; GTX632269) was used at 1:500 dilution; Goat-anti-mouse IRDye 800CW secondary antibody (licor; 926-32210) was used at 1:10,000 dilution. Goat-anti-rabbit IRDye 800CW secondary antibody (licor; 926-32211) was used at 1:10,000 dilution in western blotting.

### Western blotting

After washing with PBS, cells were lysed with 2 × SDS loading buffer (100 mM Tris-Cl [pH 6.8], 4% SDS, 0.2% bromophenol blue, 20% glycerol, 10% 2-mercaptoethanol) and then boiled for 5 min. Proteins were separated by SDS-PAGE and transferred to a nitrocellulose membrane. The membranes were incubated with blocking buffer (PBS, 5% milk, 0.05% Tween) for 1 h and then with primary antibody diluted in the blocking buffer. After three washes with PBST (PBS, 0.05% Tween), the membranes were incubated with secondary antibody. After three washes with PBST, the membrane was visualized by Western Lightning Plus-ECL substrate (PerkinElmer, NEL10500) or by Odyssey CLx Imaging System.

### *In-vitro* ligation

BsaI digested fragment were gel purified by using Gel Extraction Kit (OMEGA) and ligated with T4 ligase (New England Biolabs) at room temperature for 1h. The ligation products were phenol/chloroform extracted, precipitated by absolute ethanol, and resuspended in nuclease-free water, quantified by determining the A260 absorbance.

### *In-vitro* transcription

BAC-based sgnCoV-sGluc plasmids or purified *in-vitro* ligated products were used as templates for the *in-vitro* transcription by mMESSAGE mMACHINE T7 Transcription Kit (Ambion) according to the manufacturer's protocol. For N mRNA production, we amplified the N coding region by PCR (sense: *GGC ACA CCC CTT TGG CTC T*; antisense: *TTT TTT TTT TTT TTT TTT TTT TTT TTT TTT TTT TTT TTT TCT AGG CCT GAG TTG AGT CAG CAC*) with phCMV-N as template. Then the purified PCR product was used as template for *in-vitro* transcription by mMESSAGE mMACHINE T7 Transcription Kit as described above. RNA was purified by RNeasy mini Elute (Qiagen) and eluted in nuclease-free water, quantified by determining the A260 absorbance.

### Transfection

Cells were seeding onto 48-well plates at a density of 7.5×10^4^ per well and then transfected with 0.3 μg *in-vitro*-transcribed RNA using a TransIT-mRNA transfection kit (Mirus) according to the manufacturer’s protocol.

### Luciferase activity

Supernatants were taken from cell medium and mixed with equal volume of 2 ×lysis buffer (Promega). Luciferase activity was measured with Renilla luciferase substrate (Promega) according to the manufacturer's protocol.

## Acknowledgements

This work was supported in part by the National Science and Technology Major Project of China (2017ZX10103009), Key Emergency Project of Shanghai Science and Technology Committee (20411950103). The funders had no role in study design, data collection and analysis, decision to publish, or preparation of the manuscript.

## Author Contributions

Conceived the study: Z Yi; conducted the study: Y Zhang, W Song, S Chen, Z Yi; Data analysis: Z Yi, Y Zhang; Manuscript draft: Y Zhang, Z Yi; Resources: Z Yuan, Z Yi

